# Use of vestibular galvanic stimulation for correction of the position and of the gaze in a flight simulator

**DOI:** 10.1101/2020.05.22.111146

**Authors:** Rosario Vega, Jorge Gordillo, Vladimir Alexandrov, Tamara Alexandrova, Enrique Soto

**Affiliations:** Instituto de Fisiología, Benemérita Universidad Autónoma de Puebla, México; Facultad de Ciencias Físico Matemáticas, Benemérita Universidad Autónoma de Puebla, México; Moscow State University, Faculty of Mechanics and Mathematics, Russia

**Keywords:** Pilot training, VOR, inner ear, flight simulator, vestibular system, transcranial electrical stimulation

## Abstract

Galvanic Vestibular Stimulation(GVS) induce the sensation of movement in subjects in flight simulators and in cosmonauts, creating a cognitive simulation of movement. The system consists of a control unit, a function generator, and a power amplifier. GVS is capable of activating the neurons of the vestibular system and inducing the sensation of movement. When applied in coordination with a flight simulation program GVS modifies the eye movement control responses, electrically activating the vestibular-ocular, vestibule-colic, and vestibule-spinal reflexes. The ultimate goal of this type of stimulation is to generate augmented reality in the pilots during training or potentially also during a flight in microgravity.

## Introduction

The Vestibular System is formed by a set of biomechanical sensors located in the inner ear of all vertebrates. The vestibular system produces the appropriate reflexes and reactions to achieve and maintain a stable position of the body, as well as to stabilize the gaze (Young, 1974). This biomechanical system is composed of three semicircular canals (lateral, posterior, and anterior), oriented almost orthogonally, and two otolithic organs (saccule and utricle). The semicircular canals (SCC) detect angular movements of the head while the otolithic organs detect linear displacements (such as normal gravitational). The vestibular system generates a set of vestibule-spinal, and vestibule-colic reflexes that contribute to maintaining a stable posture, vestibule-ocular reflexes related to the maintenance of visual stability and vestibule-autonomic reflexes related to vaso-vagal stability (Fife, 2010; Cullen, 2012).

Extraocular muscles are the effectors of the vestibule-ocular reflexes (VOR). These muscles contract or relax in such a way that when receiving a neuronal input, they act to move the eyes in defined directions and in a coordinated way. It has been established that there are relationships between the SCC planes and the direction of the induced movement of the eyes and of the head (Baloh and Honrubia, 2010; Cox and Jeffrey, 2008). The activation or deactivation of the SCCs by a mechanical (or galvanic) stimulus determines, according to the SCC activated or inhibited the direction of eye movement. The eye movement system is analogous to a pulley system and by which the degree of activation of each SCC determines the direction of eye movement (CITA). The purpose of the Galvanic Vestibular Stimulation (GVS) is to stimulate the SCC of a subject to help stabilize the gaze on the indicated objective (Reynolds and Osler, 2012). Then, for example, by having a right turn in the frontal plane of a person (and stimulating on the right side), it is expected that the anterior and posterior SCC of the right side will be activated while the vertical SCC on the left side are deactivated.

### Galvanic Vestibular Stimulation (GVS)

GVS is a non-invasive method that depending on the characteristics of the stimulus (timing of stimulation, placement of the electrodes, amount of current and waveform of the stimulus), produces specific postural responses related to the activation of SCC (sensations of angular movement) and of the otolithic organs (linear displacement).

GVS produces a vestibular response without exciting other sensory inputs (Pliego, 2013; Vega et al., 2016). When an alternating current is applied at a low frequency, the stimulus has an influence on the stabilization of the gaze, and the displacement of the eyes. On the other hand, when direct current is applied, what is generated is mostly a sensation of displacement with an inclination of the body (Hlavacka et al., 1996). GVS modulates the discharge of vestibular afferent neurons. The cathodic current increases the frequency of discharge of neurons, while the anodic current decreases it (Soto et al, 2017). Cathodic or anodic GVS affects the discharge rate of afferent neurons from the semicircular canals similarly to an ipsilateral angular acceleration.

Flight simulators create a virtual reality through a dynamic simulation of the visual and auditory environment, and the use of movement platforms mimicking the airplane’s flight attitude. The present work studied the use of GVS in a flight simulator environment to define whether or not GVS may contribute to gaze stabilization.

## Material and methods

All GVS experiments were carried out taking care of the welfare of the voluntary subjects. The norms established in the Declaration of Helsinki (World Medical Association Declaration of Helsinki 2013) and the Official Mexican Standard (NOM-012-SSA3-2012) for experimentation with humans were followed. Informed consent was signed and the clinical history of each voluntary subject was made.

Voluntary subjects selected for the GVS experiments were between 18 and 30 years old (healthy without any pathology). Square pulses of 2 mA were applied; The electrical stimulation was injected through 1 cm diameter chlorinated silver electrodes (3M, Red Dot) the cathode placed in the right mastoid process and the anode centered on the forehead in the right side of the subject. Bipolar unilateral GVS was applied, which consisted of applying a constant current of 2 mA for 8 seconds. The placement of the electrodes was done assuring subjects did not show any discomfort.

The subjects sat in the cockpit of a dynamic flight simulator based on a Stewart platform. The platform as a generator of angular movements is part of a dynamic flight simulator that contains a cockpit and a display screen (Figure 1). In the experiment, the pilot’s seat, inside the cabin. The simulator movement was produced by specific algorithms to mimic the flight of an aircraft. The flight path consists of a maneuver used by pilots to evade obstacles or change their course, known as coordinated turn, which consists in making the plane perform a right or left turn in the flight simulation, changing its course, then turn again in the opposite direction to the first turn, to establish and maintain a new direction. Coordinated turn is considered, as in the case of this experiment, when the warping angle, ϕ, is non-zero (to make the rotation of the aircraft in the simulated flight environment), the angle of attack is practically zero, α ≈ 0, and the sliding angle (skidding) is zero, β = 0, as well as the speed and altitude remain constant.

**Figure 1.**
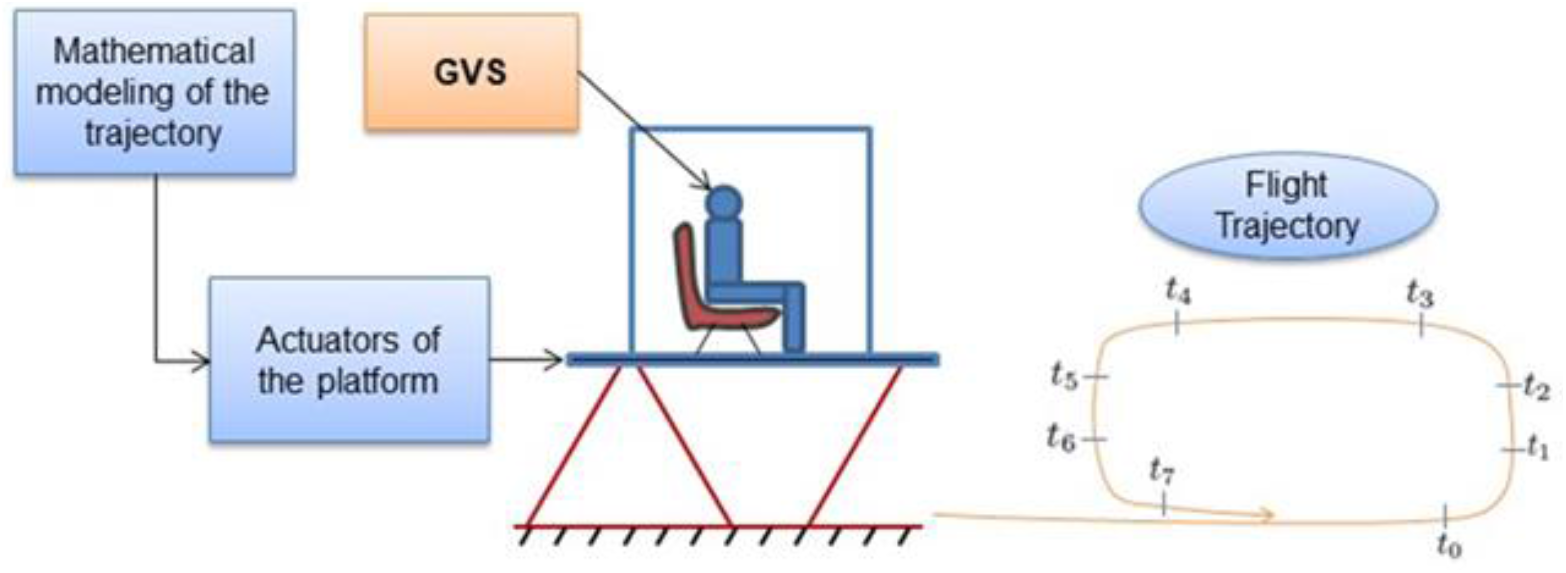
GVS and Experimental setup. Scheme depicts the experimental setup. The subject seat in the Stewart’s platform, and is subject to a dynamical flight simulation program. GVS will be applied during 10 s in two of the turns (marked), and false GVS in other two of the turns of a coordinated flight.

Linear and angular movements of the head and eye of the pilot were recorded using an ICS Video Head Impulse (Otometrics, Natus Med Inc. Taastrup Denmark). The equipment consists of a high-resolution micro camera, micro gyroscopes, and micro accelerometers all mounted on glasses, which the subject can use comfortably. This device allows us to measure with high precision the ocular movements and the movement of the head of a subject. In the case of eye movements, the reference system is located inside the Ocular Video Camera (CVO), so that, at all times, the horizontal and vertical deviations of the right eye are measured, regardless of the orientation of the subject’s head, we can also obtain their speeds and accelerations by deriving them numerically.

## Results

Three voluntary subjects participated in the execution of the experiments, and with each of them the right turn maneuver was executed three times.

Results related to the movement of the eyes and the head of the pilot (obtained with the video-ocular recording camera and gyro sensors and accelerometers). The pilot seeks to follow the movement of the platform in an appropriate manner according to his perception (response to mechanical stimuli and activation of vestibule-colic, and vestibule-ocular reflexes). The movement of the Stewart platform is not very fast, however, that generates in the pilot a feeling that he will continue with his right-turn movement, but he finds that the Stewart platform reaches its limit to then return to the starting position (0 °).

For the purpose of the experiments, GVS was used to counteract the influence of mechanical stimulation introduced by the Stewart platform. Each subject was subjected to two periods of stimulation and a period of control (platform turn without GVS). The subject seeks to follow the roll motion of the PS in an appropriate manner according to its perception (response to mechanical movement). The right turn mainly activates the right anterior SCC and inhibits the left posterior SCC, the eye moves upwards. (Fitzpatrick and Day, 2004). When EGV is applied, it is possible to stabilize the gaze close to horizontal (0 °) and manage to maintain it during all the remaining time of the experiment. In the experiment where EGV is applied, it is possible to verify that EGV inhibits the effect of the mechanical movement generated by the platform. The values for the path traveled in pre-stimulation and stimulation, where, PreEGV = 0.3366 and EGV = 1.2719 (Figure 2). It is worth mentioning that in this set of experiments, the tests were limited, however, these results show that GVS can contribute stabilizing the gaze of the subjects in the flight simulator

**Figure 2.**
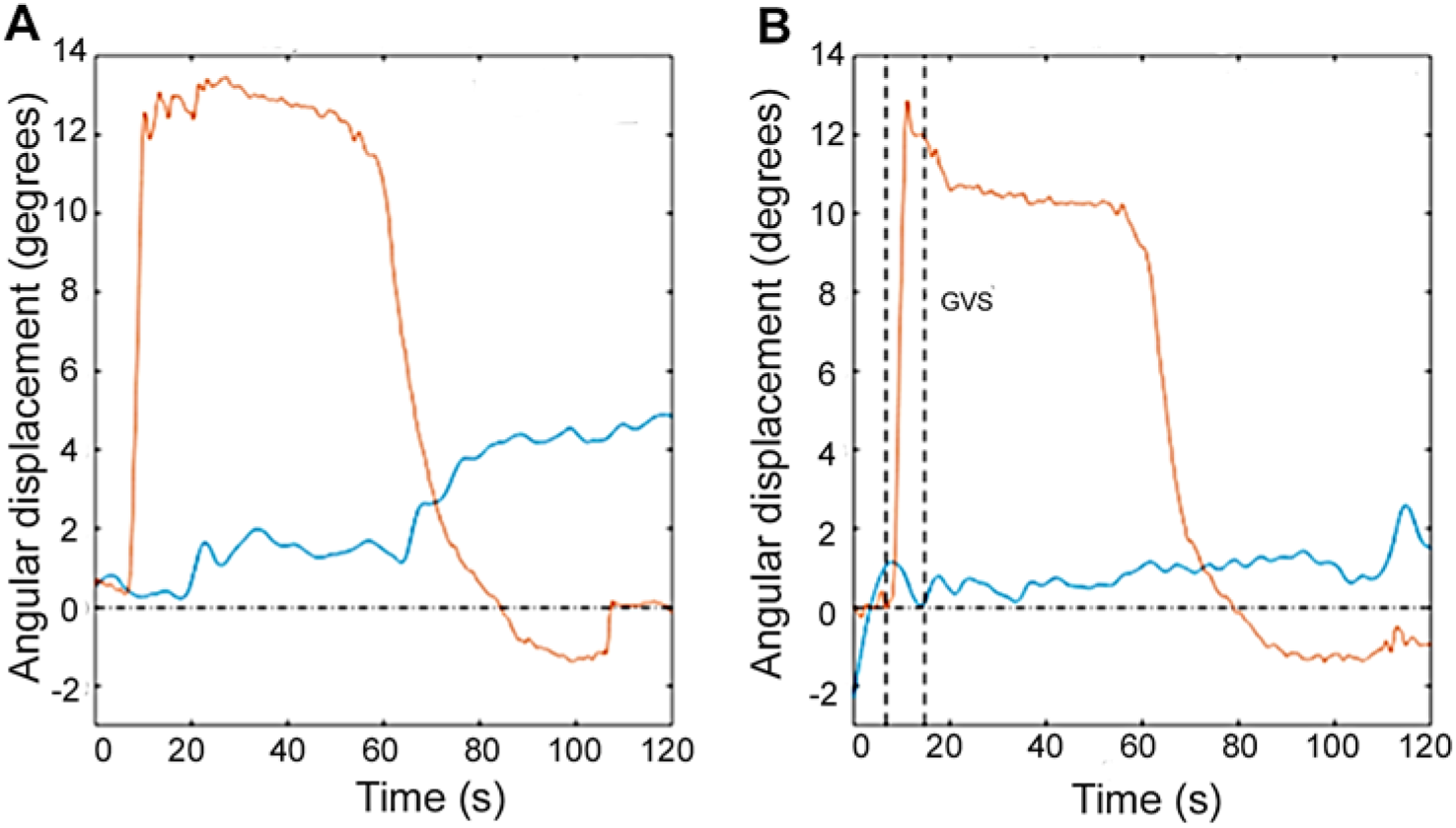
Recording of the movement of the Stewart platform and that of the eyes during the coordinated turn experiments. The roll movement of the Stewart platform was 10◦ (solid line), the right eye movement in the vertical axis of the camera (red line). In A, in control conditions. The gaze fixation error is up to 6◦ with respect to the vertical axis of the camera. In B, gaze fixation is improved by a factor of three by the use of GVS (2 μA) at the beginning of turn during 8 s.

## Discussion

We address the GVS capability to modulate the sensation of movement in a virtual reality environment in flight simulators used for the training of aircraft pilots. For pilot training, screens are used where it is projected in a flying environment, added to the use of Stewart platforms, which depend on actuators, whether electric or mechanical; the actuators increase or decrease their length to simulate the inclinations in the planes of roll, pitch, and yaw directions. In the experimental stage, we found that it is not feasible in the said platform to make turns of more than 30 ° in the plane of the yaw, and the speeds were very small due to the physical limitations of the actuators of the platform. In addition to the characteristics described above, the Stewart platform has the limitation of generating low-speed inclinations, mainly in the Z plane (yaw inclination), due to the characteristics of the actuators that generate the movements. What increases the potential of our proposal to use GVS as an aid in virtual reality or complex environments to modulate spatial orientation, whether test pilots or in microgravity.

In our results when the cathode is located on the mastoid bone of the left side of the head, it is necessary to consider a pair of vertical SCC, consisting of the right posterior and the right anterior canals, which have positive projections of the angular acceleration vector at the beginning of the turn. Mechanical action in this case leads to the activation of the right anterior canal and the absence of activation of the left posterior canal, which are a functional pair of vertical canals. Thus, the lack of activation can be replaced by a galvanic simulation of mechanical action, which leads to the activation of the corresponding afferent neurons and thus, in accordance with the contraction of the upright lower muscle of the eyeball, a moment of force occurs, which in total leads to an equality (approximate) of two moments of force (Figure 3). That is, to the absence of mechanical action on the right eyebal consequently leads to the correction of the accuracy of the gaze stabilization. Thus, the results obtained when the cathode is located on the right or left side of the mastoid bone are qualitatively the same, but the processes of galvanic stimulation are different. This property applies only to vertical semicircular canals.

**Figure 3.**
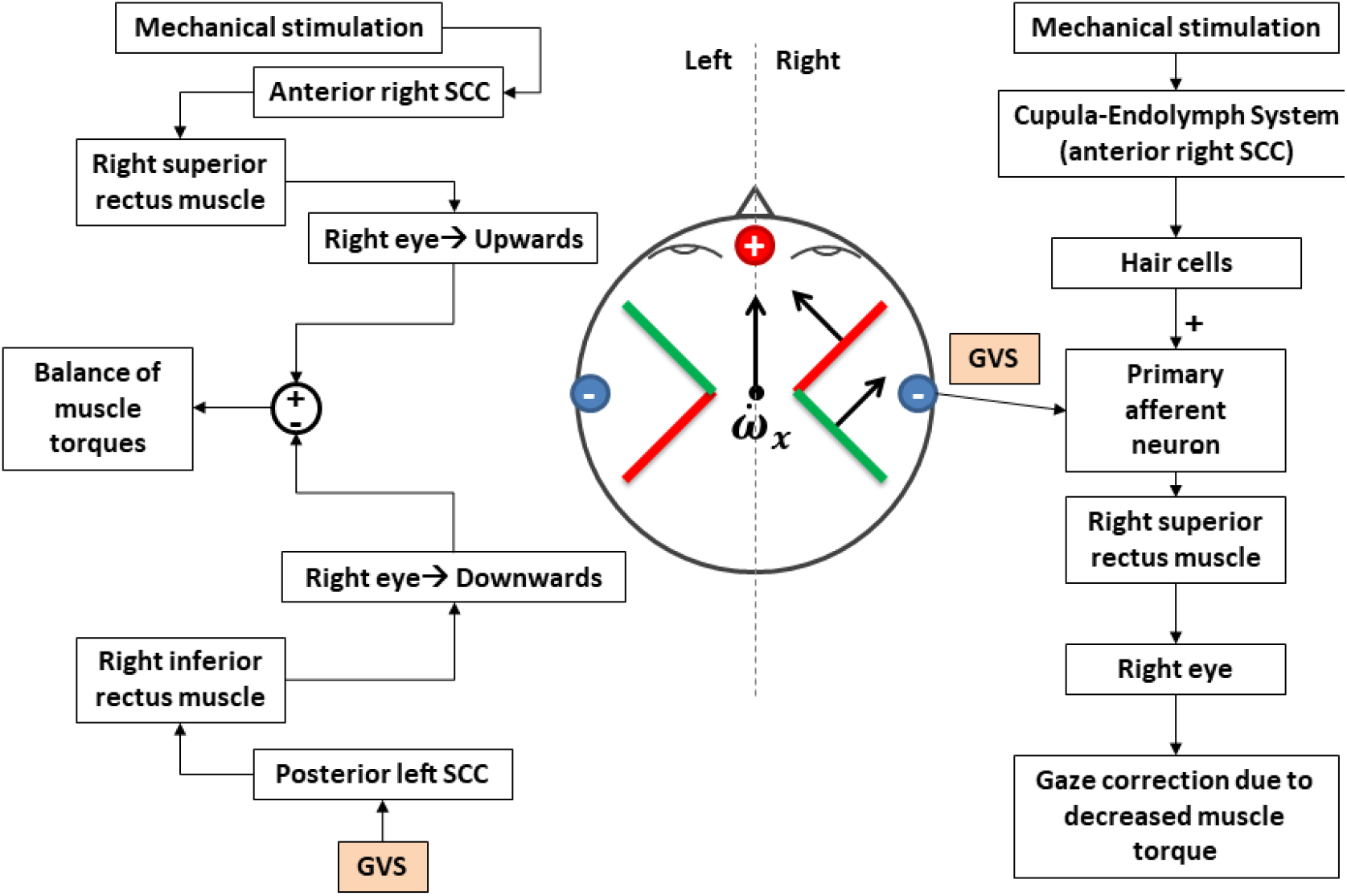
Scheme based on Ewald’s laws, showing to the left the path leading to ocular movement. To the left the path followed by GVS stimuli. In the center anterior and posterior SCC are illustrated. Showing the activation and inhibition produced by the GVS and subsequent counteraction of movement response and imporvement of gaze stabilization.

Since GVS directly stimulates reflex pathways (vestibule-ocular and vestibule-spinal), it does not require learning by users to interpret the stimulus. In addition, its prolonged use does not produce adaptation, so it will have the same effect on users regardless of how many times the device has been previously used. Our proposal of use of GVS for the training of pilots, would cover the needs for simulation in movement platforms, but not limited to, because it would be a methodology, applicable to enhance the information perceived by the pilots in training in flying simulators. Not only is the physical stimulation generated by the movements of the platform but, added to the screens and auditory signals, the vestibular device will deliver GVS to directly stimulate the organs of the balance of the pilots (Vega et al. 2016).

There are proposals in the literature to use the GVS as a tool to generate an augmented reality of movement in subjects. One proposal is that of Aoyama, which, by placing electrodes on the atrial periphery, generates realistic acceleration sensations. This method was called: walk with GVS (GVS RIDE in English); The idea is to use both the GVS and augmented reality viewers and games (Aoyama et al., 2015; Aoyama 2017).

Currently there are other methods of vestibular stimulation, one of them is transcranial magnetic stimulation, which is still under investigation. From the investigations carried out, magnetic stimulation results have been obtained ranging from dizziness to sensation of rotation and eye movement, according to the power of the applied magnetic flow. It is not yet known exactly how the magnetic flux influences, in the vestibular system. Some studies indicate that Lorentz’s strength or magnetic field strength deflects the ionic currents of the hair cells, creating the sensation of rotation. If this force is strong enough, it can cause eye movements. In cases where the subject has vestibular damage, these eye movements do not occur (Boegle and Cols., 2016).

Previous results of our research group demonstrated that GVS modulates postural responses in normal subjects (Pliego et al., 2019). The GVS can be used for the training of aircraft pilots, due to its influence on the vestibular system and gaze stabilization. The GVS can also be used for training and to improve the execution and stability of cosmonauts in orbital flight in microgravity. It is proposed to make use of the GVS to counteract the mechanical effects (unwanted displacement) of the eyeballs, to stabilize the gaze during the flight.

## Acknowledgements

This work was financed by Agencia Espacial Mexicana (AEM) and Consejo Nacional de Ciencia y Tecnología (CONACyT) grant AEM-2016-1-275058 to Rosario Vega.

